# Temperate phages increase antibiotic effectiveness in a *Caenorhabditis elegans* infection model

**DOI:** 10.1101/2024.10.28.620739

**Authors:** Rabia Fatima, Gayatri Nair, Jordan Mayol, Lesley T. MacNeil, Alexander P. Hynes

## Abstract

The bactericidal nature of obligately lytic bacterial viruses (phages) is of increasing interest for the treatment of drug-resistant bacterial infections, either administered alone or in combination with antibiotics. In contrast, temperate phages are largely ignored in a therapeutic context due to their ability to lie dormant within the bacterial host. However, these phages often undergo a lytic cycle. Furthermore, even in their dormant state they can be a considerable burden to the bacterium – most famously by their ability to switch to lytic replication in response to environmental triggers, such as antibiotics, that stress the bacterial host. Recent reports of antibiotics synergizing with temperate phages *in vitro*, termed “temperate phage antibiotic synergy” (tPAS), present a potentially scalable opportunity to make use of these abundant entities for the treatment of bacterial infections. Here we employ *Caenorhabditis elegans* as a robust *in vivo* animal model for testing the efficacy of temperate phages as adjuvants to antibiotics. *In vivo*, temperate phage Hali alone results in 60% dormancy in the bacterial survivors. However, the antibiotic ciprofloxacin can abolish dormancy of temperate phage Hali - infecting a ciprofloxacin resistant *Pseudomonas aeruginosa* clinical strain. The phage Hali-ciprofloxacin pairing increased the lifespan of *P. aeruginosa* infected worms to that of the uninfected control, at doses where neither the phage nor the antibiotic had any effect alone. Complete rescue was also observed in worms infected with a phage-carrying strain treated with the otherwise ineffective antibiotic, supporting that the phage - even in its dormant form - can greatly enhance antibiotic effectiveness. This illustrates potential “accidental” phage therapy when antibiotics are prescribed. Our work establishes *C. elegans* as a suitable model for studying the *in vivo* efficacy of tPAS and is the first *in vivo* demonstration of this synergy, greatly expanding the therapeutic potential of temperate phages.

## Introduction

In light of the growing antimicrobial resistance crisis, bacteriophages (phages) are increasingly being explored as adjuvants or alternatives to antibiotics. Most of this work has focused on a particular type of phage called virulent or strictly lytic, given their immediate bactericidal nature (1–12). However, it is estimated that 50% - 75% of all bacteria contain at least one dormant phage, termed temperate or lysogenic (13, 14). Much of the resistance to the use of temperate phages in therapy comes from concerns of further spread of fitness-benefitting genes within the bacterial population, especially the spread of toxins or antibiotic resistance cassettes. While on average 1-5 dormant phages were predicted in the genomes of per clinical isolate of *Pseudomonas aeruginosa* from cystic fibrosis patients, they were not associated with antibiotic resistance of the bacteria (15). Estimates of phage-encoded antibiotic resistance genes also appears to be overestimated due to inherent biases in bioinformatic based virome studies, with matches to low similarity and non-antibiotic resistance associated proteins (16). Of the four, out of total 421 predicted phage-encoded antibiotic resistance genes, verified in lab, none conferred antibiotic resistance (16). In addition, temperate phages are not inherently better transducers than virulent phages, many of which are capable of generalized transduction (17–19).

A key concern specific to temperate phages as treatments in the clinic comes at the level of the integrated phage (prophage) conferring to its host immunity against subsequent phage infections through a variety of mechanisms (20, 21). This leads to a rapid regrowth of phage-generated resistant ‘mutants’ and could readily contribute to treatment failure. However, carrying non-self genome such as a prophage, especially if it does not confer direct fitness benefit, can potentially pose a heavy metabolic burden on the bacteria. Lysogens of several *Pseudomonas aeruginosa* phages demonstrate defects in bacterial motility, swarming and twitching (22). As a result, the prophage can have lasting influence on their host physiology, virulence, and pathogenesis, which could be exploited for the treatment of bacterial infections.

One of the most common physiological changes in the prophage-carrying lysogen is the one from which it derives its name; the prophage can be highly responsive to environmental cues that stress their bacterial host in a process referred to as induction (23). In the well-characterized phage model Lambda infecting *Escherichia coli*, the phage can sense host DNA damage to switch from lysogenic to lytic replication, resulting in cell lysis (24). In the gastrointestinal tract of monoxenic mice, 1 to 2% of Lambda lysogenic bacteria were lysed *per* generation due to spontaneous prophage induction (25). Hence, prophage excision itself likely presents a fitness burden to the bacteria, especially in environments of high stress. Lysogens can exhibit increased sensitivity to antibiotics, monitored as decreases in minimum inhibitory concentration (MIC), compared to the parent strain (26–28). Temperate phages of *Burkholderia cepacia* (29), *E. coli* (28, 30), and *P. aeruginosa* (27) can also function synergistically with antibiotics, resulting in increases in phage production (29) or a decrease in the functionally effective dose of the antibiotic (28). When the mechanism of action specifically impacts phage lysis-lysogeny decision, this is referred to as temperate phage-antibiotic synergy (tPAS). Given that temperate phages are dominant within the human body (31), there is a strong case in favour of their therapeutic use – there existing interaction with antibiotics likely already plays a role in treatment outcomes.

There are currently several popular animal models for studying efficacy of phage therapy. Specifically for temperate phages, *Burkholderia* phage AP3 was able to significantly increase survival of wax moth larvae (15% survival at 72 h compared to 0% for untreated at 48 h) (32). In a hamster model, a four-phage cocktail targeting *Clostridioides difficile* reduced the colonization load and delayed the onset of symptoms (33). More recently, temperate phage of *Acinetobacter baumannii*, vB_AbaM_ABMM1, was able to reduce mortality in a zebrafish model (34). While useful, these models present several scalability concerns, each animal need to be individually monitored, handled and experimentally in the lab, which is time consuming and resource intensive.

In this study, we develop the nematode *Caenorhabditis elegans* as an infection model for studying the *in vivo* efficacy of a temperate phage-antibiotic combination against the multidrug-resistant bacteria *P. aeruginosa*. Due to its short and measurable lifespan and the ability to generate large, germ-free age-synchronized populations in the lab, *C. elegans* is a well-established model for the study of bacterial pathogenesis (35). Many human pathogens, including *P. aeruginosa* (36) are also pathogenic to *C. elegans*. Accordingly, *C. elegans* are also commonly employed as an *in vivo* model for studying the efficacy of antibacterial drugs (37–39). Pathogenesis in *C. elegans* of *P. aeruginosa* has been studied in three different contexts. Agar-based methods are differentiated by fast and slow killing, dependent on media composition (40). Fast killing occurs within hours, does not require live bacteria, and is mediated by bacterial secreted toxic phenazines (40, 41). In contrast, slow killing occurs over the course of several days with active intestinal colonization being a key determinant (40). A third *P. aeruginosa-* mediated *C. elegans* killing method is observed in liquid, with the same formulation as that of slow killing medium (SKM) (42). While not as well understood, this is independent of phenazine production or intestinal colonization, however, is reported to be mediated by *P. aeruginosa* secreted siderophore pyoverdine which induces a hypoxic response and subsequent worm death (42).

Only a few studies have used *C. elegans* to investigate phage therapy, focusing on virulent phages of *B. pseudomallei* (43), *Salmonella* Enteritidis (44), *Staphylococcus aureus(45), E. coli* (46), *Klebsiella pneumoniae* (46), *Enterobacter cloacae* (46), with only one study specifically for *P. aeruginosa* (47). The primary outcome measured in these studies was the ability of phages to reduce bacterial load and/or increase survival post infection. However, none of these studies tested antibiotics alone or in combination with their phage of interest. Combining temperate phages with antibiotics has demonstrated promise in previous *in vitro* studies, however *in vitro* results are not always sufficient indicators of *in vivo* efficacy (48, 49), especially in cases of antibiotic potency which can be highly sensitive to media composition, pathogen, and host factors (50, 51). As a result, here we employ a simple animal model to be able to quickly iterate and screening a large set of temperate phage-antibiotic pairings that synergized *in vitro* against *P. aeruginosa* (27). Results/Discussion

### Validation of C. elegans as an infection model

To investigate if co-administration of temperate phage and antibiotic can result in bacterial killing in an *in vivo C. elegans* model, we first validated *P. aeruginosa* infection with a clinical strain of interest, C0400. C0400 is a ciprofloxacin resistant strain where co-administration of temperate phage Hali and ciprofloxacin was able to re-sensitize the strain to the antibiotic *in vitro* (27). The strain PA14 is commonly used in *C. elegans* for studying *P. aeruginosa* pathogenesis (40, 41) and served as our control. For our experiments, we employed worms carrying a temperature-sensitive mutation (*bn2*) in the germ-line proliferation regulator *glp-4*. Mutant animals maintained at the restrictive temperature of 25°C are sterile, which bypasses the need to separate adults from their offspring in survival assays (52). In slow killing assays, dependent on host colonization, both *P. aeruginosa* strains colonized the intestine at levels higher than *E. coli* OP50 control, with no differences between PA14 and C0400 (Fig. S1a). Similarly, *P. aeruginosa* fed worms maintained on SKM begin dying after three days, much faster than OP50 control, with complete population death observed by day 10 (Fig. S1b). The lifespan data closely matches previously reported survival data of the *glp-4(bn2)* strain exposed to OP50 and PA14 (52), validating that the clinical *P. aeruginosa* strain C0400 is suitable for investigating tPAS in this *in vivo* model.

**Fig 1.**
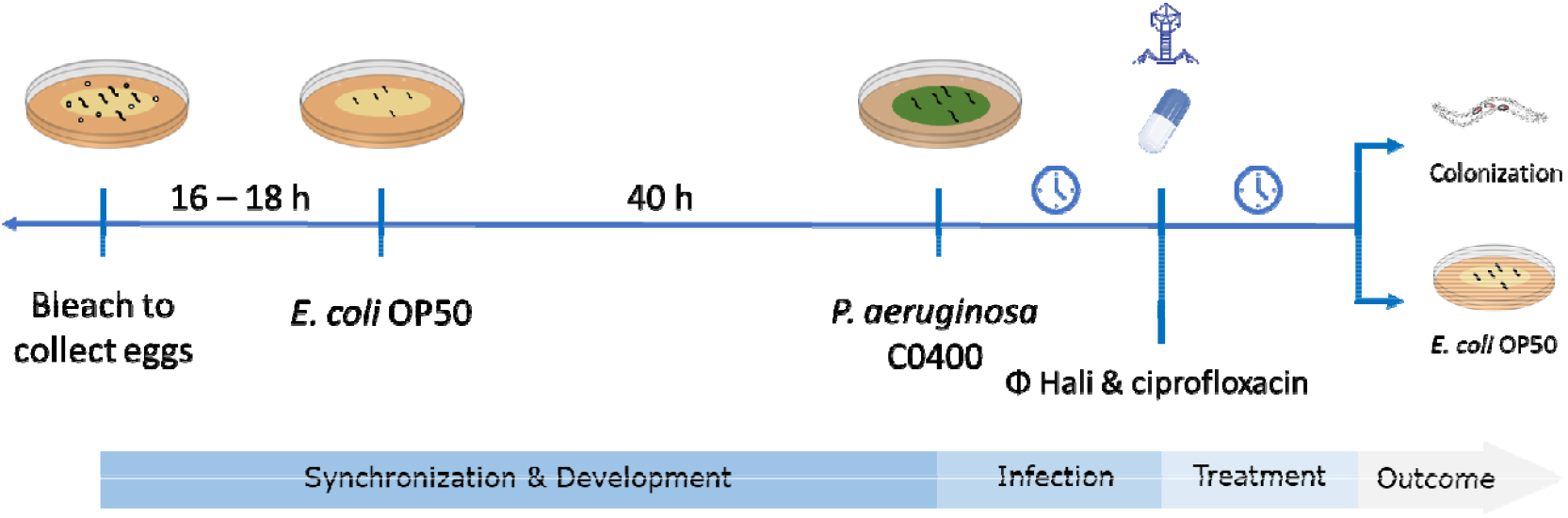
Representation of the assay used for testing efficacy of temperate PAS in a *C. elegans* model of *P. aeruginosa* infection. Worms are bleached to collects eggs which are hatched and synchronized to L1 stage. Animals are developed on *E. coli* OP50 and infected with *P. aeruginosa*, followed by phage and/or antibiotic treatment. Outcome of treatment is monitored by bacterial colonization and lifespan on regular bacterial diet. Each assay was done in independent biological replicates, with technical triplicates plated for assessing bacterial load.

### Ciprofloxacin selects against lysogens in vivo

To test whether tPAS can reduce *P. aeruginosa* loads *in vivo*, we used two different treatment durations, 4 h and 18 h, after infection on SKM (Fig. 2). We treated worms with ciprofloxacin across a range of antibiotic concentrations (0.25 μg/mL – 180 μg/mL), alone and in combination with phage Hali. For 4 h treatment, there was no effect observed at both higher and lower dose (Fig. S2a-b). After 18 h of treatment, we observed a high degree of biological variability between replicates at lower concentrations of ciprofloxacin (Fig. S2c-d). There was no reduction in colonization when treated with phage alone and only 1-2 log_10_ reduction observed with 100 μg/mL ciprofloxacin (Fig. 2a). Similarly, there was no synergistic reduction in bacterial load for tPAS for either treatment duration.

**Fig 2.**
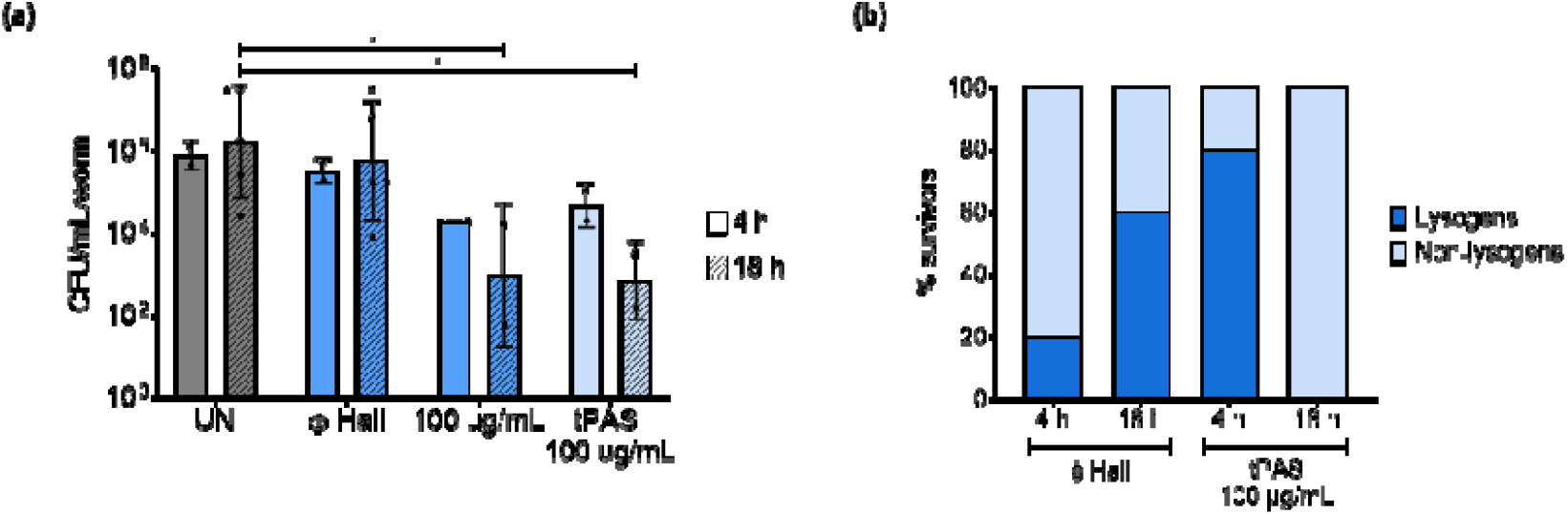
Phage Hali and ciprofloxacin selects against lysogen survivors *in vivo*. **a)** *C. elegans* intestinal colonization measured as log10 cfu/mL/worm after 48 h C0400 infection followed by 4 h or 18 h treatment with phage Hali (1 × 10^9^ pfu/mL) ± ciprofloxacin. Data shown as mean ± SD, each data point i a biological replicate plated in technical triplicates, except ones denoted by triangle, which are single technical replicate, with of ∼50-100 worms per condition. The two treatment durations were tested in independent trials. Values were compared using two-way ANOVA with a multiple comparison Tukey test post-hoc with conditions only compared within their own treatment duration, *P-value 0.01-0.05. **(b)** Frequency of lysogeny within survivors of phage Hali alone and phage Hali + ciprofloxacin (tPAS). Twenty survivors were tested for each condition.

Despite no reduction in bacterial loads observed, it was still possible that tPAS biased the survivor profile, selecting against lysogens. Treatment with phage alone resulted in an increase in lysogens from 20% to 60% as treatment duration increased (Fig. 2b). This regrowth of lysogens is expected as they are immune to further phage infection due to superinfection immunity and aligns very closely with our previous report that phage Hali results in 75% lysogens after an overnight challenge *in vitro* (27). In comparison, tPAS resulted in primarily lysogen population within 4 h treatment (80% compared to 20% for phage alone) and a reduction in frequency of lysogeny to zero with longer treatment duration. Compared to administering phage by itself, tPAS selectively biases the phage away from lysogeny *in vivo*, completely preventing the regrowth of lysogens that would otherwise form 60% of the population. This formation of lysogens (4 h) that are then depleted is consistent with the previously reported mechanism of this interaction to be operating through induction (27).

In short, while all treatments had – at best – modest effects on the bacterial load, combining phage and antibiotic drastically altered the nature of those bacteria, selecting against lysogens. We sought to determine whether this bias was enough to rescue the worms after treatment.

### Combined temperate phage and ciprofloxacin rescues P. aeruginosa infected worms

To test if selecting for a different survivor population would be enough to rescue the worms, we carried out the same assay but moved the animals to their standard *E. coli* OP50 diet after 18 h treatment to measure lifespan. We performed the lifespan assay with a shorter 4 h infection as we did not observe any rescue from even the antibiotic at high doses with a longer infection duration, suggesting that worms were moribund (data not shown). With a shorter infection, when left untreated, the worms died within maximum thirteen days compared to eighteen plus days for the uninfected OP50 control as they reach the end of their lifespan (Fig. 3). Treatment with phage Hali alone was not only ineffective, but it also showed a worse survival outcome compared to untreated controls in two of the three trials (Fig. 3a). Ciprofloxacin alone at low antibiotic dose of 2 μg/mL did not reliably rescue survival alone or in combination with phage (Fig. S3a). At a much higher antibiotic dose, 92 μg/mL, the antibiotic alone was sufficient to rescue (Fig. S3b). However, we could not reliably discern the effect of tPAS as both antibiotic alone and tPAS survival curve matched the uninfected OP50 control in two trials.

**Fig 3.**
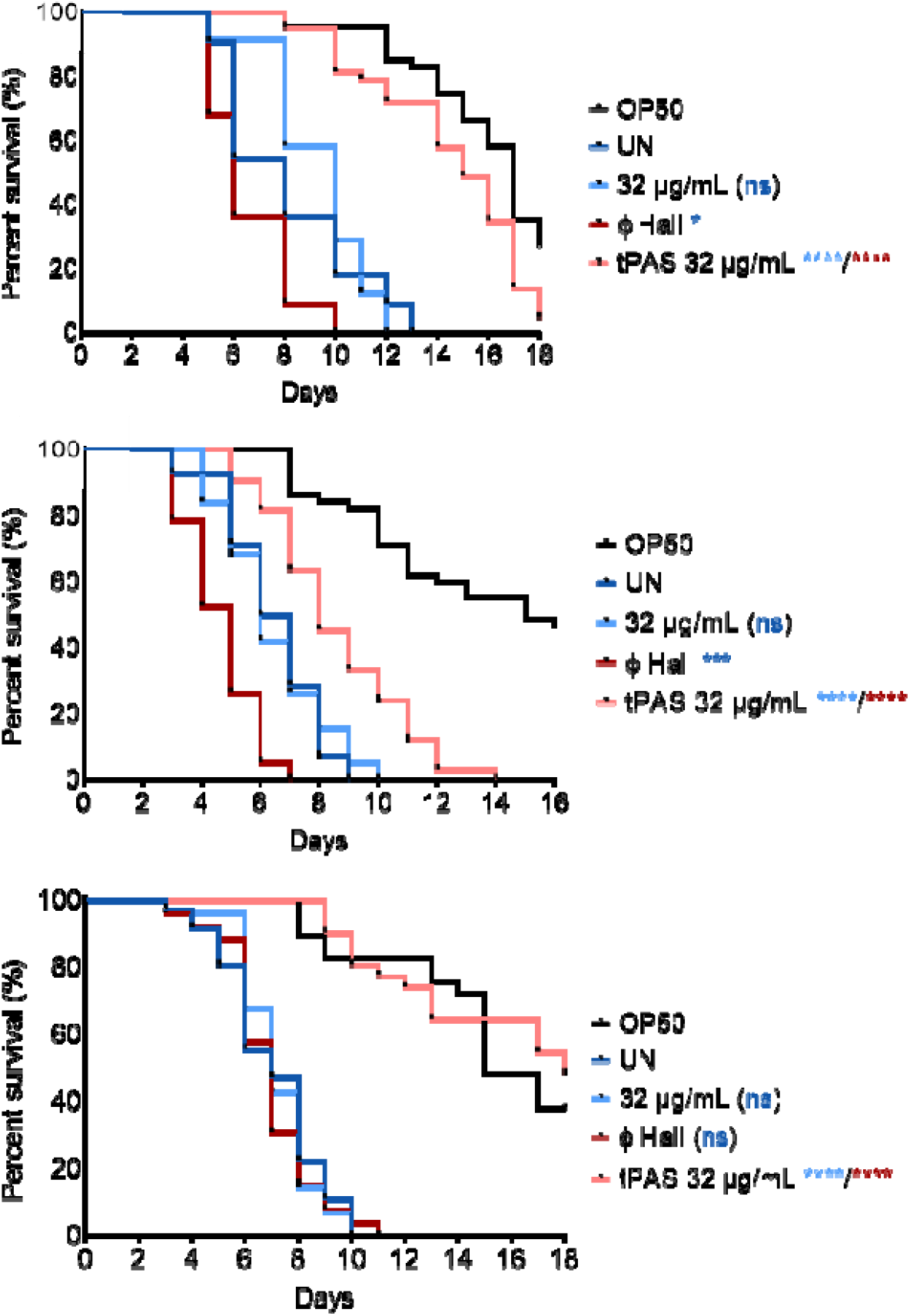
Phage Hali and low dose ciprofloxacin increase survival of *P. aeruginosa* infected worms on standard *E. coli* OP50 diet after treatment. Life span of worms on OP50 after 4 h *P. aeruginosa* infection followed by 18 h phage Hali ±ciprofloxacin treatment. Approximately 50 worms were added to each condition at the time of treatment, tested in three independent biological replicates performed on separate days. Survival curves were compared using the Log-rank (Mantel Cox) test with one pair compared at a time, ns = not significant, * P ≤ 0.05, *** P ≤ 0.001, and **** P ≤ 0.0001. Phage alone and antibiotic alone were compared to the untreated, and tPAS to the phage alone and antibiotic alone, as denoted by colour of the significance value.

We were primarily interested in antibiotic concentrations that were lower than the effective dose and were therefore insufficient to increase survival. Ciprofloxacin treatment alone at 32 μg/mL did not increase survival (Fig. 3). In comparison, tPAS had a marked improvement in lifespan to levels comparable to the uninfected OP50 control in all trials, compared to either treatment alone. In short, the *in vitro* work that had shown synergy between phage Hali and ciprofloxacin translated to a robust, reproducible *in vivo* increase in worm survival at concentrations where neither agent alone had any detectable effect.

We asked whether survival was driven by bacterial load, performing the colonization assays as carried out earlier (Fig 2) but found that treatment outcomes for these shorter 4 h infections did not correlate with bacterial loads (Fig S4). For instance, 32 μg/mL ciprofloxacin with or without phage were both successful in eradicating *Pseudomonas*, even though the antibiotic-alone treatment had no effect on lifespan (Fig. S4). The most likely explanation is that the phage-antibiotic combination is benefiting the worms through either faster eradication, qualitatively different endpoints for the bacterium (phage-mediated lysis rather than antibiotic-mediated death), or that the presence of transient lysogeny is in some way mitigates the virulence of the strain.

### Lysogen infected worms can be rescued with antibiotic

To determine whether the formed lysogens were differentially virulent, we infected worms with wild type C0400 and a C0400 phage Hali lysogen, followed by ciprofloxacin treatment and subsequently monitoring survival on the standard *E. coli* OP50 diet. Without treatment, the pathogenicity of the lysogen and wild type strains did not differ (Fig. 4a). Intriguingly, lysogen-infected worms could be completely rescued with 32 μg/mL ciprofloxacin, which was insufficient to rescue worms infected with the wild type C0400. This strongly supports the notion that the tPAS observed by co-administration of the phage and antibiotic is primarily driven by the interaction between the prophage and the antibiotic, as any phages induced (spontaneously or by the antibiotic) would have little effect on the lysogens, which are immune to superinfection.

**Fig 4.**
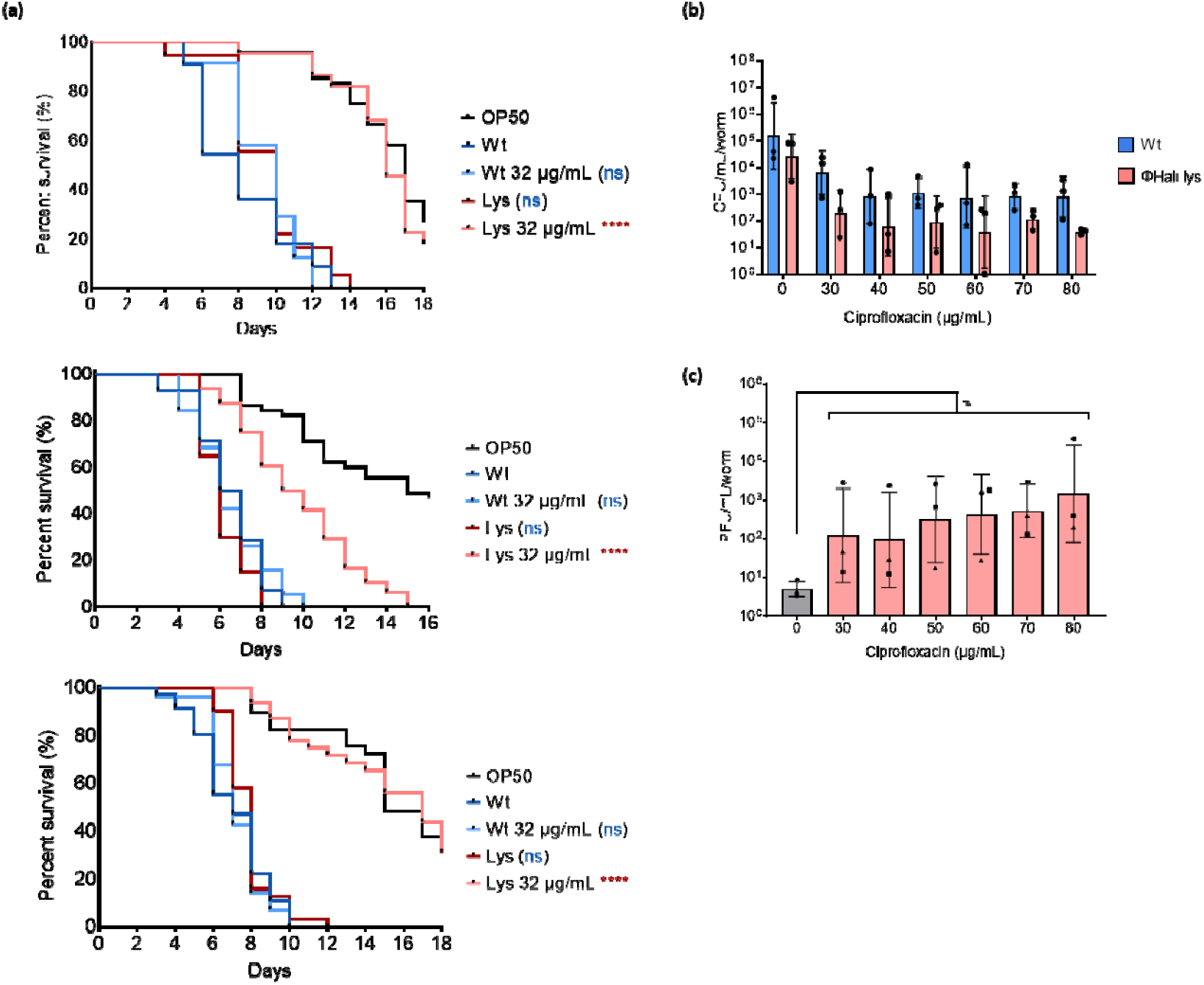
Phage Hali lysogen infected worms can be rescued with antibiotic alone but the lysogen is not more sensitive to the antibiotic. **(a)** Representative life span of worms on OP50 after 4 h *P. aeruginosa* C0400 wild type and phage Hali lysogen infection followed by 18 h ciprofloxacin treatment. Approximately 50 worms were added to each condition at the time of treatment, tested in three independent biological replicates performed on separate days. Survival curves were compared using th Log-rank (Mantel Cox) test with one pair compared at a time, ns = not significant, * P ≤ 0.05, *** P ≤ 0.001, and **** P ≤ 0.0001. Lysogen untreated was compared to wild type untreated, and antibiotic treatment was compared to its respective no antibiotic control, as denoted by colour of the significance value. **(b)** *C. elegans* intestinal C0400 wild type and phage Hali lysogen colonization measured as CFU/mL/worm after 48 h infection followed by 18 h treatment with ciprofloxacin. Bacterial load wa compared using two-way ANOVA with a Bonferroni test. All comparisons were not significant. **(c)** Corresponding phage quantification shows as PFU/mL/worm for the lysogen infected, and ciprofloxacin treated worms. Phage titer compared using a one-way ANOVA with a Dunnett test, ns represent not significant. All bar graph data shown as mean SD where each data point is a biological replicate carried out with roughly 100 worms per condition, plated in technical triplicates, except for ones denoted b circle and square in (c) which were plated in single replicates.

This finding is particularly important mechanistically, because one possible explanation for tPAS would be that the phage administration results in a modest reduction of the bacterial load prior to the regrowth of lysogens, and that at this lower bacterial density the bacteria are more sensitive to the antibiotic – a concept known as density-dependent antibiotic sensitivity (53, 54). However, here the phage and antibiotic are truly synergistically interacting. This similar rescue observed to tPAS (Fig. 3) also means that irrespective of whether the phage is co-administered or if it exists as a prophage, it can increase the effectiveness of the antibiotic if it is antibiotic inducible.

Considering that lysogens are often more sensitive to environmental stressors because of carrying prophages that are responsive to these triggers (26–28), we assessed if carrying a prophage substantially burdens the bacteria *in vivo*, especially in the presence of antibiotic, by measuring intestinal bacterial colonization of worms infected with wild type or lysogen and treated with ciprofloxacin. Similar to the earlier colonization results reported in Fig S2c-d, we observed a high degree of biological variability in colonization post treatment with no observed difference between wild type or lysogen colonization post ciprofloxacin treatment (Fig. 4b). In short, in our colonization assays we could often see eradication without rescue, and in our rescue assays, we could see rescue without difference in colonization. This reinforcing the idea that bacterial loads at the times selected are not a good predictor of treatment efficacy, but the kinetics of their death (or the manner of it – phage mediated vs antibiotic mediated) can drastically impact survival. It is worth noting that, while not statistically significant, there was a 10-100-fold higher phage count inside the lysogen infected worms after antibiotic treatment, consistent with induction (Fig. 4c), and given the trend towards reduced bacterial loads (Fig. 4b) this likely underestimates the magnitude of induction. These data are consistent with *in vitro* report that phage Hali lysogens shows 4-fold lower ciprofloxacin MIC compared to wild type, and that this is due to prophage induction (27).

While we were unable to correlate increases in worm survival to reduction in bacterial load, the increased survival of phage Hali lysogen-infected worms treated with ciprofloxacin taken together greater phage counts observed from the lysogen infected worms suggests that carrying an inducible phage could impose a major fitness cost under mild antibiotic stress. Given the abundance of prophages within bacterial hosts (14, 31) and that pathogens are well-known reservoirs of prophages (55, 56), this effect is likely already taking place during antibiotic treatments, in a form of “accidental phage therapy”. It is important to note that this is likely phage dependent, not just with regards to its integration site and receptor, but especially in its ability to respond to antibiotic triggers. The *P. aeruginosa* LES strain common in cystic fibrosis patients is known to contain several prophages that can be induced with ciprofloxacin *in vitro* (57). In addition, from patient data, the induction of these phages was shown to play a crucial role in regulation of the pathogen in CF lungs (58). However, it is challenging to correlate the effects of long-term antibiotic prescription to pathogen load and phage induction since the data are incomplete. However, James et al. (2014) do propose that treatments that result in phage induction could be promising approaching to tackling *P. aeruginosa* load in these patients, not without caution in case of phage encoded virulence factors (58).

## Conclusion

In phage therapy, temperate phages are avoided due to their ability to lysogenize. However, these have an enormous untapped potential due to their ability to respond to host stressing environmental cues, such as antibiotics, to switch into lytic replication, killing its host in the process. Here we show that *C. elegans* is a suitable model for studying *in vivo* efficacy of tPAS, but this can easily be extended to traditional PAS with virulent phages. While we did not observe synergistic reduction in bacterial load, ciprofloxacin strongly selects against phage Hali lysogens. In addition, even if lysogens do arise, they can be more sensitive to the antibiotic *in vivo*, compared to the parental strain. While phage Hali is completely insufficient to rescue the worms, phage and ciprofloxacin combined increased the lifespan of *P. aeruginosa* infected worms to levels comparable to the uninfected control, at an otherwise ineffective antibiotic dose. Complete rescue can be seen for the lysogen-infected worms treated with the antibiotic. The phage, even in its prophage form, greatly enhanced antibiotic effectiveness. This suggests to us that this kind of ‘accidental’ phage therapy already often occurs when antibiotics are prescribed.

## Supporting information

Raw data for all figures

## Acknowledgement

The authors would like to Mercedes DiBernardo for helping set up the worm strains and sharing her expertise on the colonization assay, as well as Sommer Chou for providing training on the worm sorter. We would also like to thank Dr. Gerry Wright for access to the *P. aeruginosa* clinical isolates from the Wright Clinical Isolate Collection at the Institute for Infectious Disease Research (McMaster University, Ontario) and Dr. Lori Burrows for PA14.

R.F was supported by an NSERC Canadian Graduate Scholarship, Master’s and Doctoral scholarship. A.P.H. acknowledges funding through the Natural Sciences and Engineering Council of Canada (NSERC) Discovery Grant 2018-05996 and the Farncombe Family Chair in Phage Biology.

R.F. jointly with A.P.H. conceived this study. R.F. performed all the assays, with help from G.N. and J.M. for bacterial and phage plating of the colonization experiments. L.M. provided the worm strains, *E. coli* OP50, and all the experiments were performed at L.M. laboratory. R.F. and A.P.H contributed to the writing of the manuscript.

## Material and Methods

### Bacterial strains and growth conditions

*P. aeruginosa* strain PA14 was obtained from Dr. Lori Burrows, McMaster University and *P. aeruginosa* C0400 was from the McMaster IIDR Wright clinical isolate collection. Bacterial strains were grown in 10 mL of lysogeny broth (LB) at 37°C with 250 rpm shaking (Ecotron, Infors HT, Quebec, Canada). For growth on solid media, 1% (w/v) of LB (lysogeny broth) agar and 0.75% (w/v) of LB soft agar was used. All plates were incubated at 37°C from overnight growth.

To prepare *C. elegans* plates, bacterial strains were streaked out on 1% LB agar plates and incubated overnight. A single colony was inoculated into 10 mL LB broth, followed by overnight incubation with shaking. To seed plates, 200 μL of overnight culture was used for 100 mm plates and 100 μL was used for 60 mm plates. The plates were incubated overnight at 37°C prior to transferring worms on to them.

### Phage propagation and titration

Phage Hali infecting *P. aeruginosa* PA14 and C0400 was used in this study. Phage lysates were prepared either using a primary or secondary amplification. Briefly, primary amplification was performed by inoculating 10 mL LB broth with bacteria and phage from frozen glycerol stocks for amplification to take place overnight. Secondary amplification was performed as needed to increase phage titer by inoculating 10 mL of same day culture prepared in LB with 50 μL of primary amplification. Phages in filtrate were quantified by standard spot test or full plate plaque assay.

Multiplicity of infection (M.O.I) was calculated based on titer and volume of bacteria and phage being combined using the following formula:
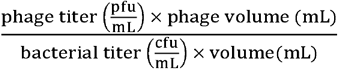. The bacterial titer was determined as previously described (27).

### *C. elegans* strains and growth conditions

All experiments were performed using *C. elegans* SS104, a temperature sensitive sterile strain, *glp-4* (*bn2*) available from the *Caenorhabditis* Genetic Center (CGC, https://cgc.umn.edu/). Worms were maintained at 15°C on 100 mm nematode growth media (NGM) plates (59) seeded with overnight cultures of *E. coli* OP50, prepared as indicated above. Eggs were collected by hypochlorite treatment (60). Eggs were rocked approximately 18 h at 20°C to synchronize to L1 larval stage. L1 worms were plated on 100 mm NGM plates seeded with *E. coli* OP50 and developed to young adults at 25°C, restrictive temperature at which they are sterile, for approximately 40 h.

### Antibiotic stock preparation

Ciprofloxacin (hydrochloride) was obtained from Cayman Chemicals (Catalog 14286-5, Ann Arbor, Michigan, USA). Working stocks were prepared in nuclease free water. The minimum inhibitory concentration of each newly prepared batch was re-evaluated prior to use as reported previously (27).

### *C. elegans P. aeruginosa* infection

For *P. aeruginosa* infection, young adults were washed off NGM OP50 plates with 10 mL M9 and washed three times to remove excess bacteria. Worms were transferred to 100 mm Slow Killing Media plates (SKM; 0.35% peptone, 50 mM NaCl, 2% agar, 1 mM CaCl_2_, 5 μg/ml cholesterol, 1 mM MgSO_4_, 20 mM KH_2_PO_4_, 5 mM K_2_HPO_4_) seeded with different *P. aeruginosa* strains for appropriate infection duration as indicated in figure legends. Worms were also plated on *E. coli* OP50 for the life span assay as a control.

### Preparing *P. aeruginosa* infected worms for treatment

After infection with *P. aeruginosa*, worms were washed off SKM plates with 10 mL M9, followed by another three M9 washes. Each wash step consisted of pelleting the worm 2 min at 1200 rpm and removing the supernatant, except for the third wash in which they were allowed to settle by gravity. To further dilute the bacteria, 1 mL of M9 with 10 ug/mL of tobramycin was added and worms were pelleted by centrifugation, followed by transfer to an unseeded 100 mm 10 μg/mL SKM tobramycin plate for 45 min in the biosafety cabinet to allow remove surface attached bacteria. After crawling, they were washed off with SKM tobramycin plates with 10 mL M9 tobramycin and subsequently washed another three times with 10 mL M9 tobramycin and final three times with M9 alone to dilute out the tobramycin.

### Colonization assay

To measure the efficacy of tPAS on bacterial colonization, approximately 50-100 infected worms were added to 2 mL tube manually, along with ciprofloxacin (final concentration tested 0.5 μg/mL – 32 μg/mL at 2-fold increments, then 32 μg/mL - 100 μg/mL in increments of 10 μg/mL) and/or phage Hali at 1 × 10^9^ pfu. The final volume was topped up to 1 mL using SKM broth. SKM alone was used a negative control. Tubes were incubated for 18 h with 150 rpm horizontal shaking and colonization assay was carried out subsequently.

Following treatment, worms were settled by centrifugation for 2 min at 1200 rpm or by gravity when working with more than 10 samples. After removing the supernatant, worms were washed three times with 1 mL M9, followed by once with 1 mL 10 μg/mL M9 tobramycin. To remove surface attached bacteria, they were transferred to 60 mm 10 μg/mL SKM tobramycin plates for 45 min. At this point, the total number of animals for each condition was counted.

Worms were washed off plates with 2 mL 120 μM levamisole in M9 (LM buffer). The number of worms remaining on the plate was counted to determine the number of animals per condition going into the lyses step. Animals were washed with 1 mL 10 μg/mL tobramycin LM buffer; once for 4 h treatment and three times for 18 h treatment. To dilute out the tobramycin, worms were additionally washed three times with 1mL M9. In the last wash, 500 μL of the supernatant was transferred into a new tube to quantify any residual bacteria, which will be deducted from bacterial load calculation later.

Approximately ten sterile silica carbide beads were added to each tube and worms were lysed for 1 min at speed 5 using the Bead Mill 4 (Fisherbrand, catalog 15340164). Lysed fractions and the last wash supernatants were serial diluted 10-folds in LB up to 10^−8^ and 3 μL was spotted onto a 1% LB plate in technical triplicates, along with one replicate for the last wash. LB only and M9 with beads were spotted as control. Plates were incubated at 32°C to prevent overgrowth and colony forming unit were counted after an overnight incubation. Bacterial load was determined by subtracting the counts from the last wash and the cfu/mL was normalized to the number of animals per condition.

### Frequency of lysogeny

Frequency of lysogeny in the survivors was determined as previously described (27). Twenty colonies for each condition were streak purified three times on 1% LB plate and inoculated in 200 – 250 μL LB broth overnight in a narrow 96 well plate. Wild type culture, C0400 phage Hali lysogen, and LB broth were added as control. The plates were incubated for 18-24 h at 37°C with no agitation. The inoculated plates were stamped on an agar overlay of wild type C0400 using a disposable pin replicator. Plates were also stamped onto 1% LB with no agar overlay to serve as growth controls. Frequency of lysogeny was calculated as percent of the total number of survivors that resulted in clearing of the wild type host.

### Survival on OP50 after treatment assay

*C. elegans* were infected with *P. aeruginosa* strains, or OP50 as control, for 4 h and washed off as described above. The volume of infected worms per strain was adjusted to 1 worm/μL and approximately 100 worms were sorted into each well of a 96 well plate using the COPAS FP (Union Biometrica, Holliston, MA). These 100 worms were then transferred to a 2 mL tube, along with 50 μL of 20x ciprofloxacin stock (final concentration tested 0.25 μg/mL – 32 μg/mL at 2-fold increments, then 32 μg/mL - 100 μg/mL in increments of 10 μg/mL), phage Hali at 1 × 10^9^ pfu. Nuclease free water was used as substitute for antibiotic and LB in place for phage. The final volume was topped up with SKM broth to 1 mL. The worms were incubated in treatment for 18 h shaking horizontally at 150 rpm.

Following treatment, animals were washed three times with 1 mL M9, allowed to settle by gravity in the third wash, and once with 1 mL 10 ug/mL M9 tobramycin. They were transferred to 60 mm SKM tobramycin plates for 45 min to crawl, after which they were washed off with 2 mL M9 tobramycin. They were washed once more with 1 mL M9 tobramycin, ending with final three washes with M9 alone.

After the last wash, animals were moved to 60 mm SKM plates seeded with OP50. Plates were incubated at 25°C and scored daily for survival starting on day three, with dead worms removed each day. Survival was plotted using the Kaplan–Meier method.

### Statistical tests

Details of the statistical tests can be found in the figure legend. N denotes a biological replicate performed from an independently grown culture with appropriate technical replicates indicated in the figure caption. Quantitative values are represented by mean ± SD. All statistical analysis was done using GraphPad Prism 8.0.2 or 10.3.0 (GraphPad Software, Inc., CA, US), with P value ≤ 0.05 is considered significant.

## Supplementary

**Fig S1.**
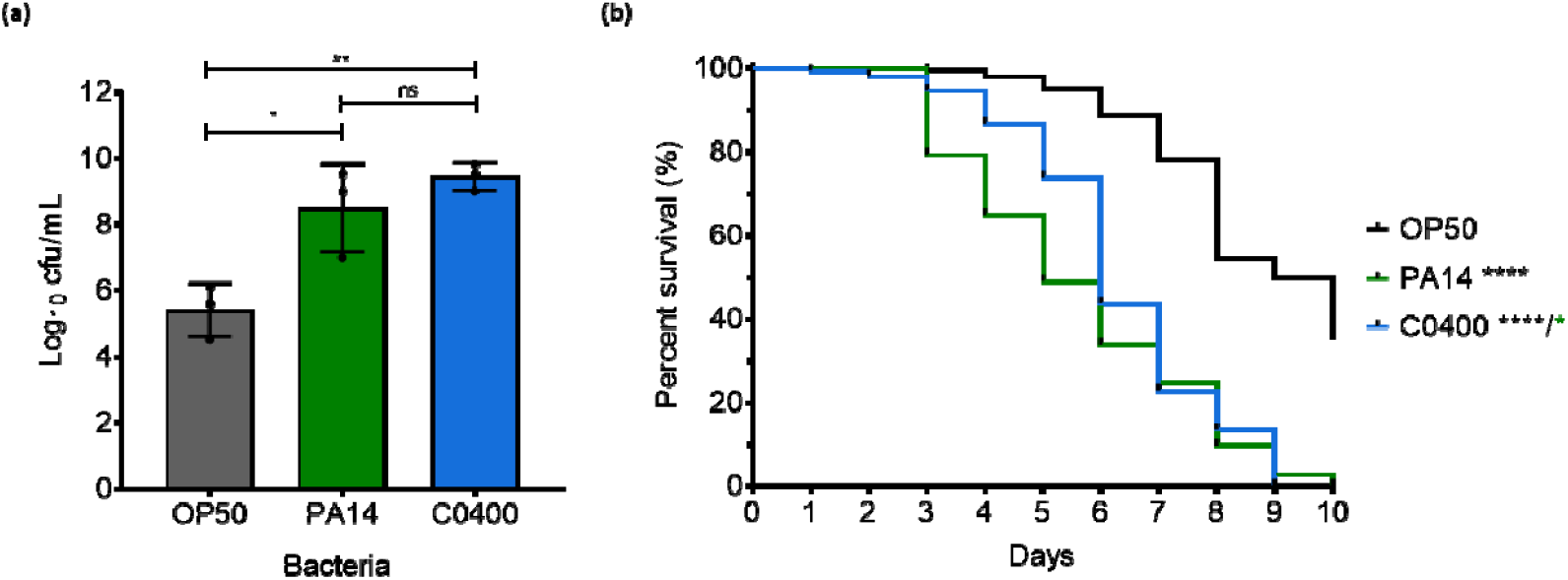
*P. aeruginosa C0400* can colonize and kill *C. elegans*. **(a)** *C. elegans* intestinal colonization measured as cfu/mL after 24 h growth on *E. coli* OP50 or *P. aeruginosa* strains (PA14 or C0400). Data shown as mean ± SD of three biological replicates, each in single technical replicate. Values were compared using a one-way ANOVA with a Tukey multiple comparison test with *P ≤ 0.05, **P ≤ 0.01, and ns representing not significant. **(b)** Representative survival curve of worms exposed to OP50, PA14, or C0400 in slow killing assay. Curves were compared using the Log-rank (Mantel Cox) test with on pair compared at a time, *P ≤ 0.05 and **** P ≤ 0.0001. Each condition is 150-200 animals. The colour of the significance value represents the comparison performed, with black represent comparison to control and green to PA14.

**Fig S2.**
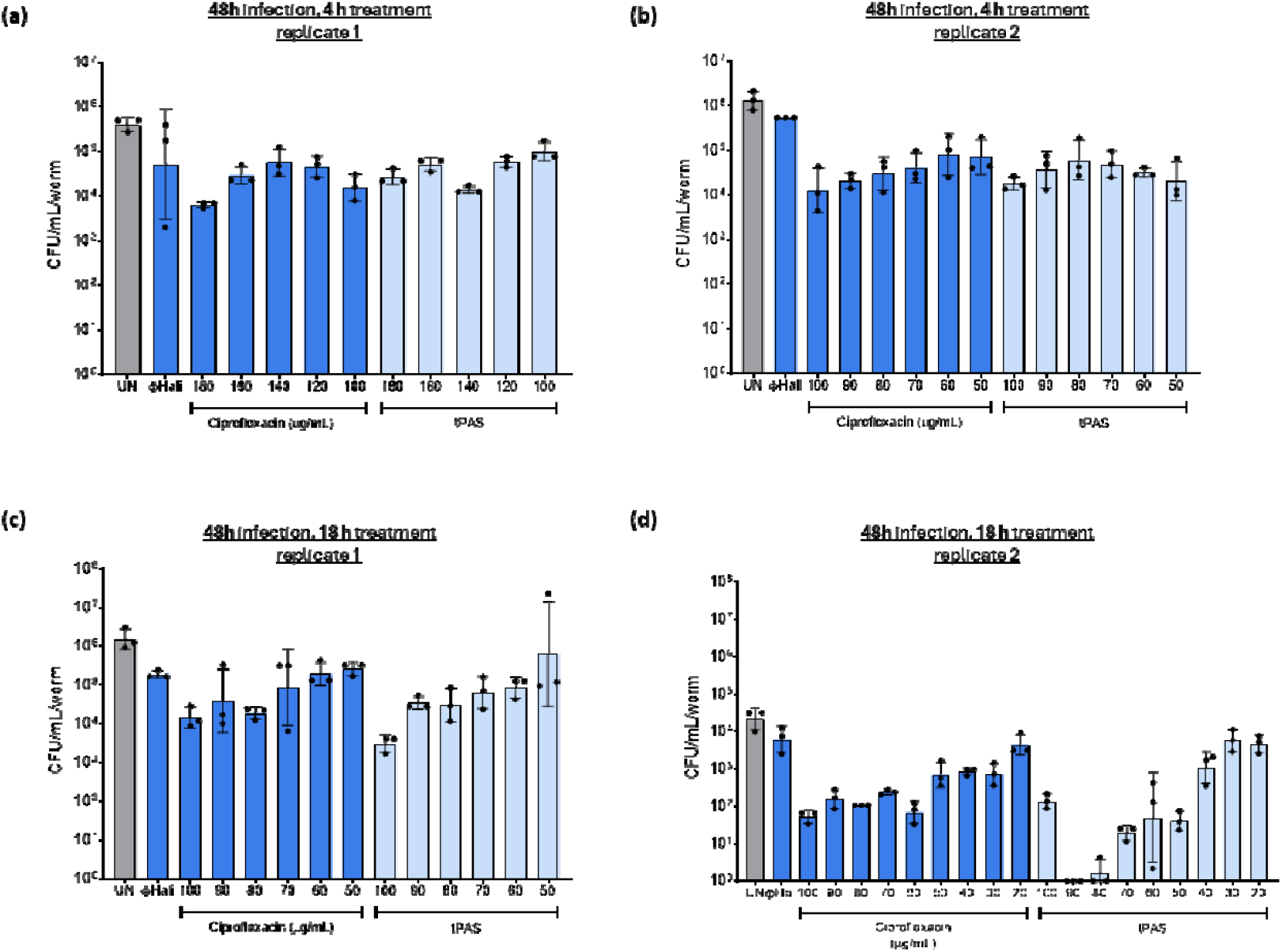
Longer treatment duration shows variability in bacterial colonization. *C. elegans* intestinal colonization measured as cfu/mL/worm after 48h C0400 infection followed by **(a-b)** 4 h **or (c-d)** 18h treatment with phage Hali (1 × 10^9^ pfu/mL) ciprofloxacin. Each graph shows a single biological replicate carried out with approximately 50-100 worms. UN denotes untreated. Data shown as mean ± SD where each data point is a technical replicate of bacterial plating.

**Fig S3.**
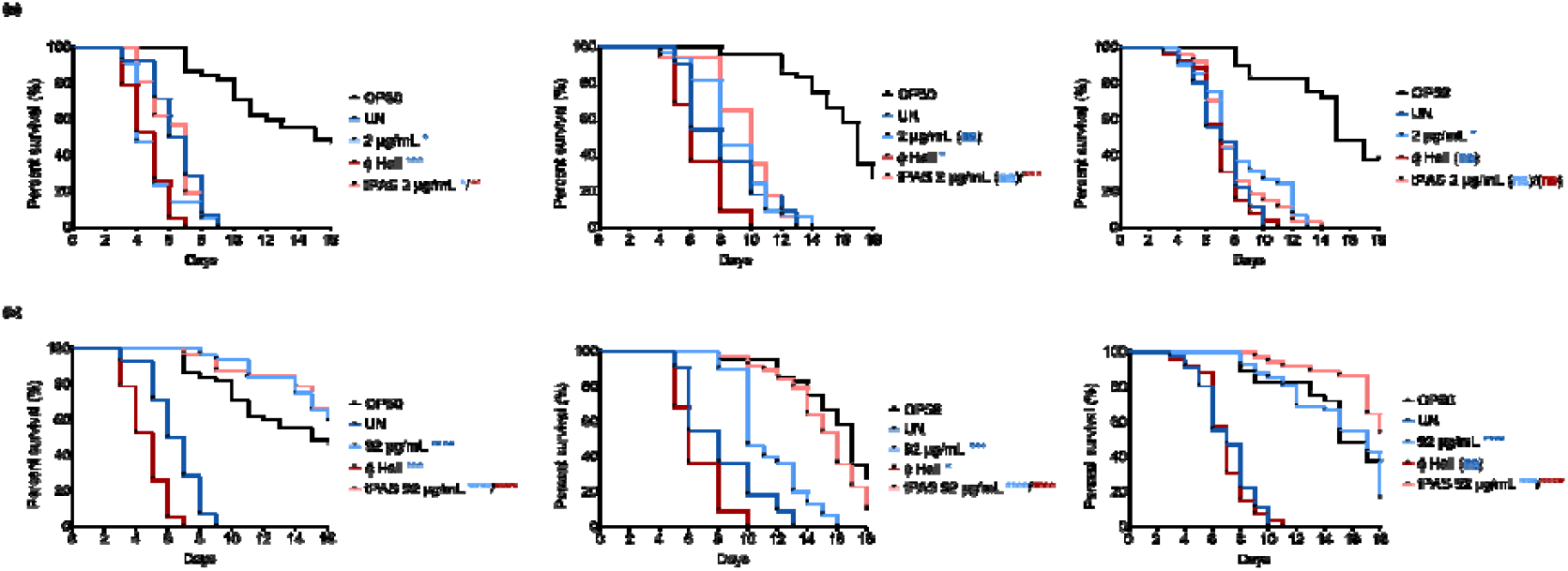
Ciprofloxacin alone at 92 ug/mL could rescue life span of *P. aeruginosa* infected worms on standard *E. coli* OP50 diet after treatment. Survival curve of worms on OP50 after 4 h *P. aeruginosa* infection followed by 18 h phage Hali ciprofloxacin treatment, **(a)** low dose 2 μg/mL and **(b)** high dose 92 μg/mL. Approximately 50 worms were added to each condition at the time of treatment. Independent biological replicates are shown for each concentration. Survival curves were compared using the Log-rank (Mantel Cox) test with one pair compared at a time, ns = not significant, * P ≤ 0.05, *** P ≤ 0.001, and **** P ≤ 0.0001. Phage alone and antibiotic alone were compared to the untreated, and tPAS to the phage alone and antibiotic alone, as denoted by colour of the significance value.

**Fig S4.**
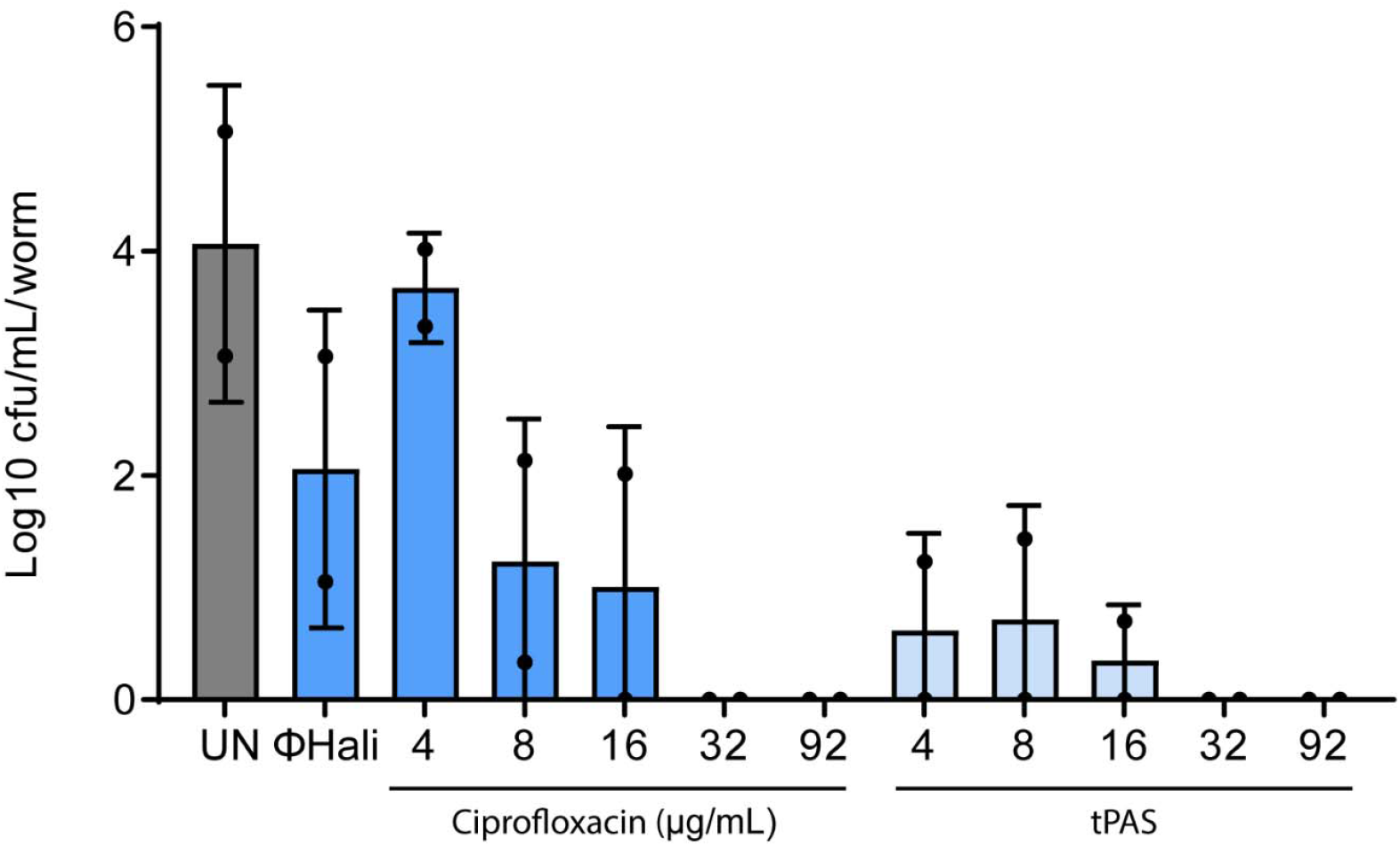
Shorter infection prior to treatment does not provide enough resolution to measure bacterial colonization. *C. elegans* intestinal colonization measured as log10 cfu/mL/worm after 4h C0400 infection followed by 18 h treatment with phage Hali (1 × 10^9^ pfu/mL) +ciprofloxacin. UN denotes untreated. Data shown as mean ± SD where each data point is a biological replicate performed in technical triplicates with ∼100 worms each.

## Notes

### Competing Interest Statement

The authors have declared no competing interest.

### Summary of Updates

Additional replicates added to colonization assays, text adjusted accordingly.

